# A chromosome-scale assembly of the sorghum genome using nanopore sequencing and optical mapping

**DOI:** 10.1101/327817

**Authors:** Stáphane Deschamps, Yun Zhang, Victor Llaca, Liang Ye, Gregory May, Haining Lin

## Abstract

The advent of long-read sequencing technologies has greatly facilitated assemblies of large eukaryotic genomes. In this paper, Oxford Nanopore sequences generated on a MinION sequencer were combined with BioNano Genomics Direct Label and Stain (DLS) optical maps to generate a chromosome-scale *de novo* assembly of the repeat-rich *Sorghum bicolor* Tx430 genome. The final hybrid assembly consists of 29 scaffolds, encompassing in most cases entire chromosome arms. It has a scaffold N_50_ value of 33.28Mbps and covers >90% of *Sorghum bicolor* expected genome length. A sequence accuracy of 99.67% was obtained in unique regions after aligning contigs against Illumina Tx430 data. Alignments showed that 99.4% of the 34,211 public gene models are present in the assembly, including 94.2% mapping end-to-end. Comparisons of the DLS optical maps against the public *Sorghum Bicolor* v3.0.1 BTx623 genome assembly suggest the presence of substantial genomic rearrangements whose origin remains to be determined.

---

Single-molecule “long-read” sequencing technologies allow complex structural aspects of genomes, such as ribosomal DNA or transposon regions, to be elucidated in a way that shorter second-generation sequencing reads typically cannot, mainly by producing long contiguous sequences from single-molecule PCR-free DNA templates (1–2). Two long-read sequencing technologies, from Pacific Biosciences (“PacBio”) and Oxford Nanopore Technologies (“ONT”), are commercially available. Oxford Nanopore sequencers measure changes in ionic current when the DNA fragments translocate through protein nanopores in a semi-synthetic insulated membrane. During translocation, nucleotides within a DNA fragments blocks the flow of ions, creating specific signals that are then translated into base calls. Because the detection process does not require enzyme-based nucleotide incorporation and detection of fluorescent signals, ONT sequencing read length is theoretically only limited by the length of the DNA fragment translocating through the pore. Consequently, sequences >1Mbps in length have been reported. This read length advantage is partially offset by the fact that individual raw sequencing reads tend to have higher error rates than second-generation sequencing reads, thus requiring sequence correction and post-assembly sequence polishing to generate high quality sequencing data. Shortly after its inception in 2014, studies have shown that the ONT technology could be used for assembling small microbial genomes (3–5). Continuous improvements in chemistry, base calling and sequencing throughputs have recently allowed researchers to use the technology for producing high-quality whole genome assemblies for larger genomes, in combination with Illumina sequencing reads to improve ONT contig sequence accuracy, in species such as human (6), *Arabidopsis thaliana* (7) and the wild tomato species *Solanum pennellii* (8).

Numerous genomic assembly projects, including maize (9) and *Aegilops tauschii* (10), have generated chromosome-scale scaffolds when combining long-read PacBio sequences with complementary long-range scaffolding technologies, such as BioNano Genomics optical maps or Hi-C proximity ligation (11). The optical mapping technology developed by Bionano Genomics has been used in genome sequencing projects to provide contig sequence validation, error correction, and high-level scaffolding resulting in significant increases in contiguity and quality (12, 13). Bionano systems enable the imaging stretching and imaging of up to megabase double-stranded DNA fragments that have been fluorescently labeled at short specific sequence motifs. Labeled fragments, or molecules, are then digitized and assembled into long-range physical maps that can be integrated into long sequence contigs. More recently, BioNano Genomics introduced a new labeling concept, called Direct Label and Stain (or “DLS”). In DLS, the Direct Labeling Enzyme 1 (DLE-1) attaches a single fluorophore to specific sequence motifs. Unlike previous Bionano labeling strategies, which were based on nicking endonucleases, DLS does not create damage in DNA at specific sites, thus avoiding loss of DNA information and truncation due to fragile sites, resulting in a significant increase in continuity. Most tested complex genomes yield 50 to 100 times higher maps N_50_ than those generated using an endonuclease approach, often leading to the assembly of full chromosome arms or even entire chromosomes.

The ability of the ONT technology combined with a long-range scaffolding technology such as optical mapping, to rapidly and cost-effectively create chromosome-scale assemblies of repeat-rich plant genomes, has not been demonstrated yet. Such assembly could be used for many applications, including the study of structural variations in large regions, expected recombination frequencies in specific regions, target sequence characterization and modification for gene editing, or genomic characterization of transgenic material.

In this study, the *Sorghum bicolor* inbred Tx430 transformation line (14) was chosen as a model plant system for whole-genome sequencing and assembly. Sorghum represents an economically important staple crop - it is the third most produced cereal crop in the United States, and its adaptability to drought and high temperatures has made it a preferred crop to grow in semi-arid regions. While its genome, at ~730Mbps, is smaller than other more complex crop genomes such as maize (~2.3Gbps) (15) or soybean (~1.2Gbps) (16), its high repeat content (~61%) and presence of large transposon structures makes it an attractive model for testing the effectiveness of the ONT chemistry for overall correctness and contiguity of a complex plant genome assembly. In addition, a reference *Sorghum bicolor* genome has been produced via Sanger sequencing for genotype BT×623 (17), making it a valuable resource for the direct evaluation of the ONT-based assembly. In order to assess the structural integrity of the Tx430 ONT assembly, contigs were aligned to a Tx430 optical map generated specifically for this study with the new Direct Label and Stain (“DLS”) technology from BioNano Genomics. The overall contiguity and accuracy of the assembly was further defined through alignments of ONT contig sequences to multiple datasets. Results shown here suggest that the ONT technology can be used to quickly and cost-effectively generate informative assemblies and, in combination with long-range DLS optical maps from BioNano Genomics, can generate chromosome-scale assemblies to assess the overall structural integrity of large and repeat-rich plant genomes.

## RESULTS

### MinION sequencing metrics and characterization

High-molecular weight genomic DNA was extracted from sorghum Tx430 adult leaf tissue using a protocol which was originally designed for the extraction of DNA for BAC library construction (18). Prior to library construction, “short-read” DNA fragments were generated after shearing DNA to ~8Kbps with a g-TUBE (Covaris), while “long-read” DNA fragments were generated after size-selecting un-sheared DNA to 20Kbps and above with a PippinHT system (Sage Science). Libraries were prepared with the ONT Ligation Sequencing Kit 1D (SQK-LSK108) and, for a selected number of “long-read” libraries, with the ONT Rapid Sequencing Kit 1D (SQK-RAD002), then sequenced for 48 hours using MinION R9.4 and R9.5 flow cells. The resulting fast5 files were processed using the Albacore base caller (version 2.0.1). A total of 4.58 million reads totaling 32.6Gbps, were generated from the “short-read” libraries, with an average read length of 7,114bps (read length N_50_ 8,039bps) while 1.9 million reads were generated from the “long-read” libraries, totaling 33.84Gbps, with an average read length of 17,459bps, a read length N_50_ of 27,204bps and a maximum read length of 335,563bps. Assuming a 732Mbps sorghum genome size, “short-read” and “long-read” sequences amounted to 90.7X equivalent of the genome.

84.23% of the raw “short reads” and 78.28% of the raw “long reads” had a Q score above the Q score passing cutoff of 7 (including 60.07% and 57.87% of the short reads and long reads, respectively, with a Q score comprised between 9 and 12). However, 68.54% of the raw “short reads” and 76.99% of the raw “long reads” had a sequence identity to the sorghum public reference genome below 80%, confirming what has been observed in previous studies, and further suggesting that the Albacore sequencing pipeline software used here was not optimized for unamplified plant genome DNA.

### MinION sequence correction and assembly

Several tools, including Canu (19) and SMARTdenovo (20), have been used for producing high-quality genome assemblies from single-molecule, error-prone, long reads. The ONT-based genome assembly of the tomato species *Solanum pennellii* indicated that a combination of Canu for correction and SMARTdenovo for assembly generated the best results (8). Canu also was described as requiring almost two orders of magnitude more CPU hours than SMARTdenovo (8). Therefore, for this study, the use of Canu was limited to sequence correction only, and SMARTdenovo was used for the *de novo* assembly of Canu-corrected reads. Only raw ONT reads larger than 2Kbps in length were submitted to Canu correction. After correction, only corrected reads larger than 5Kbps were submitted to SMARTdenovo assembly. After assembly, ONT contigs were polished twice with Pilon (21), using an Illumina Tx430 whole-genome shotgun dataset produced from a paired-end library with an average fragment size of ~400bps. A total of 691.64MM Illumina 150bps reads were produced, totaling 103.7Gbps, or 141X sequencing coverage of the ~732Mbps sorghum genome. Results from the assembly are summarized in Table 1. As shown, the Tx430 genome assembly produced a total of 734 contigs, with a contig N_50_ of ~3Mbps and a mean contig length of 899.9Kbps. The assembly contained 660.5Mbps. The largest contig was 16.1Mbps in length. Two rounds of Pilon polishing with the Illumina Tx430 whole-genome shotgun dataset slightly decreased the number of contigs to 723, while increasing the size of the assembly to ~671.8Mbps (Table 1). Distribution of contig lengths (Supplementary Figure 1) indicates that 97% of the assembly, or 651.3Mbps, is contained within the largest 400 contigs.

**Table 1:** Summary of ONT assembly metrics. (Unit = bps) Assembly metrics are shown before polishing with Pilon, after one round of polishing with Pilon (1X) and after two rounds of polishing with Pilon (2X). Contigs generated after two rounds of polishing were used for subsequent analysis.

|  | CANU +<br>SMARTdenovo | CANU +<br>SMARTdenovo +<br>Pilon (1X) | CANU +<br>SMARTdenovo +<br>Pilon (2X) |
| --- | --- | --- | --- |
| Number of Contigs | 734 | 724 | 723 |
| Total Length | 660,557,552 | 671,283,418 | 671,866,842 |
| Average Contig Length | 899,942 | 927,187 | 929,276 |
| Minimum Contig Length | 8,556 | 8,623 | 8,655 |
| Maximum Contig Length | 16,114,705 | 16,329,467 | 16,337,099 |
| N<sub>25</sub> Contig Length | 7,217,337 | 7,344,315 | 7,351,898 |
| N<sub>50</sub> Contig Length | 3,005,621 | 3,051,816 | 3,053,182 |
| N<sub>75</sub> Contig Length | 1,248,066 | 1,261,559 | 1,254,124 |

### Initial assessment of accuracy and completeness

To estimate the overall accuracy of the ONT assembly, contig sequences were aligned against the public v3.0.1 *Sorghum bicolor* assembly. Alignments were first performed after splitting contigs into 10Kbps sequences and mapping them to the public assembly using minimap2 (22). Results then were filtered for the best hit for every 10Kbps sequences with a coverage cutoff of 50%. This approach has the advantage of performing end-to-end alignments of 10Kbps sequences, rather than computing sequence identities from gapped fragments. The average identity for all best hits to the public assembly was 95.95%. Since the source material used to generate the v3.0.1 *Sorghum bicolor* assembly was from a different genotype (BTx623), the observed average sequence identity could be related to natural variations between the two genotypes, or the fact that some regions in the public assembly, including centromeres, were not fully sequenced. To better predict ONT error rates and determine the sequence accuracy of the Tx430 ONT contig sequences, the Illumina Tx430 whole genome shotgun data used in this study for Pilon polishing were mapped against the ONT contigs using BOWTIE2 (23). After alignment, ONT sequence accuracy was determined against the uniquely aligned Illumina reads. Results showed 92.94% of uniquely aligned Illumina reads (corresponding to 229.88MM reads) exhibited >99% sequence identity (including 84.95% with 100% sequence identity) to the ONT assembly while only 81.03% of the reads had >99% sequence identity (64.78% with 100% sequence identity) to the public v3.0.1 assembly. The resulting average accuracy rate of all ONT contigs to the uniquely aligned Illumina reads was 99.62%, with the remaining discrepancies after two rounds of Pilon polishing dominated by mismatches. Additional polishing of the ONT contig sequences with nanopolish (3), followed by two additional rounds of Pilon polishing, only slightly increased the overall accuracy to the uniquely aligned Illumina reads to 99.67%, while decreasing the overall alignment rate of Illumina data from 97.74% to 96.84%. Therefore, only the initial ONT contigs polished twice with Pilon were used for subsequent analysis.

To test the overall completeness of the Tx430 ONT assembly, contig sequences were compared to individual chromosome sequences from the public v3.0.1 *Sorghum bicolor* genome assembly. NUCmer v3.1 (MUMmer 3.23 package) (24) generated a total of 2,376 partial contig alignments with sequence identity to the reference averaging 97.67% and ranging from 95.02% to 100%. Comparisons were made using MUMmerplot 3.5 (MUMmer 3.23 package) on the NUCmer alignments, which were subsequently filtered for 1-on-1 alignments and rearrangements with a 20Kbps length cutoff (Figure 1). Despite the presence of random nucleotide stretches in centromeres and other regions of the v3.0.1 *Sorghum bicolor* assembly, the results indicated a high genomic completeness rate of the ONT assembly, with sufficient sequence homology to recover contigs covering most of the v3.0.1 *Sorghum bicolor* assembly.

**Figure 1:**
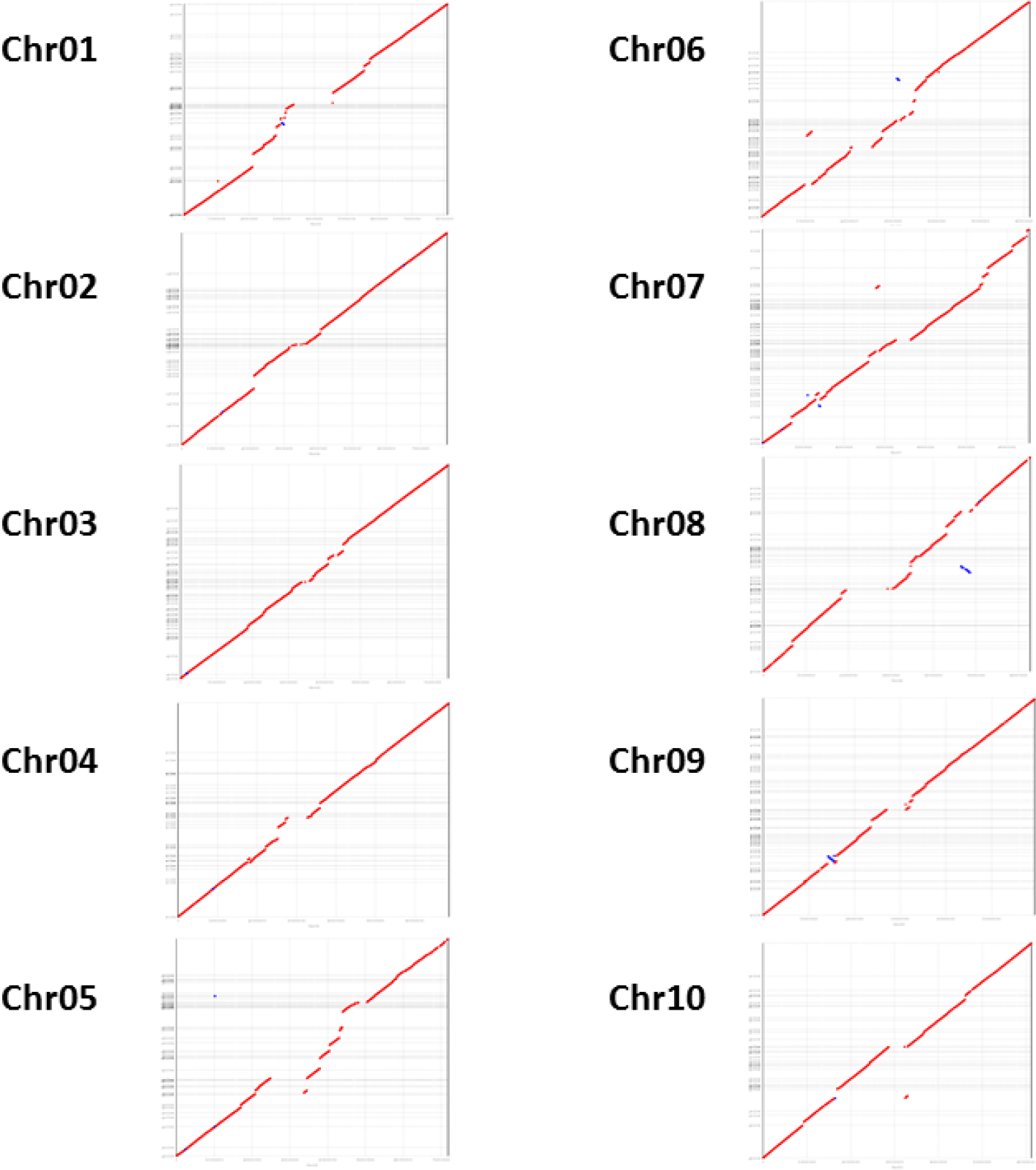
MUMmerplot comparison of Tx430 ONT assembly (Y-axis) with all 10 chromosomes from the public *Sorghum bicolor* v3.0.1 genome assembly (X-axis). ONT contigs were aligned to the BTx623 assembly with NUCmer and results were subsequently filtered for 1-on-1 alignment and rearrangements with a 20Kbps length cutoff. BTx623 chromosomes are labeled by number and contain multiple megabase-scale regions in the form of unresolved nucleotide sequences.

Finally, in order to assess the overall genic completeness and accuracy of the assembly, the 34,211 gene models from the public *Sorghum bicolor* v3.0.1 release (corresponding to a mixture of *bona fide* protein-coding genes and a small number of *ab initio* predictions based on EST support from other crop species) were mapped using BOWTIE2 to the 724 contigs generated in this study. A total of 32,217 gene models, or 94.2%, were found to be entirely mapped to at least one ONT contig, with a mean sequence identity of 98.77%. An additional 1,794 transcripts (5.2%) were mapped partially to at least one contig, while only 200 transcripts were absent from the assembly. Such result indicates a genic completeness level very much on par with the *Solanum pennellii* ONT-assembly study (genic completeness of 94.86%).

One of the potential applications of an ONT whole-genome assembly is the systematic determination of genomic target sites for CRISPR/Cas9 editing in specific transformation lines, such as sorghum Tx430. One example is the centromere-specific histone H3 (Sb-CENH3) gene (25). The Tx430 genomic region encompassing the Sb-CENH3 gene was determined via sequencing and *de novo* assembly of a Tx430 library prepared with the Chromium platform from 10X Genomics, and sequenced on an Illumina sequencer. The Sb-CENH3 gene was found, overlapping on two 10X Genomics contigs, 6,426bps and 16,686bps in size, respectively. Alignment of the 6,426-bp and 16,686-bp contigs with the ONT assembly showed full coverage and sequence identities of 99.9% and 99.8%, respectively, suggesting that the ONT assembly alone is suitable for detecting and characterizing potential CRISPR/Cas9 target sites in the sorghum Tx430 transformation line.

### Generation of a BioNano DLS contig map and comparison to the public genome assembly

To generate chromosome-level maps in sorghum Tx430, a Bionano Saphyr system was used in combination with the recently released Direct Label and Stain (DLS) technology. To construct optical maps in sorghum Tx430, a total of 1,224,604 DNA fragments with lengths ranging from 150Kbps to 2,706Kbps and a N_50_ length of 286Kbps were imaged and digitized (Table 2). The final *de novo* assembly yielded 79 maps, with a total combined length of 732.1Mbps. The resulting Map N_50_ was 33.78Mbps, with the largest map being 47.64Mbps. Out of 79 maps, 32 maps accounted for 99.5% of the total assembly aligned to the BTx623 reference (Figure 2).

**Table 2:**
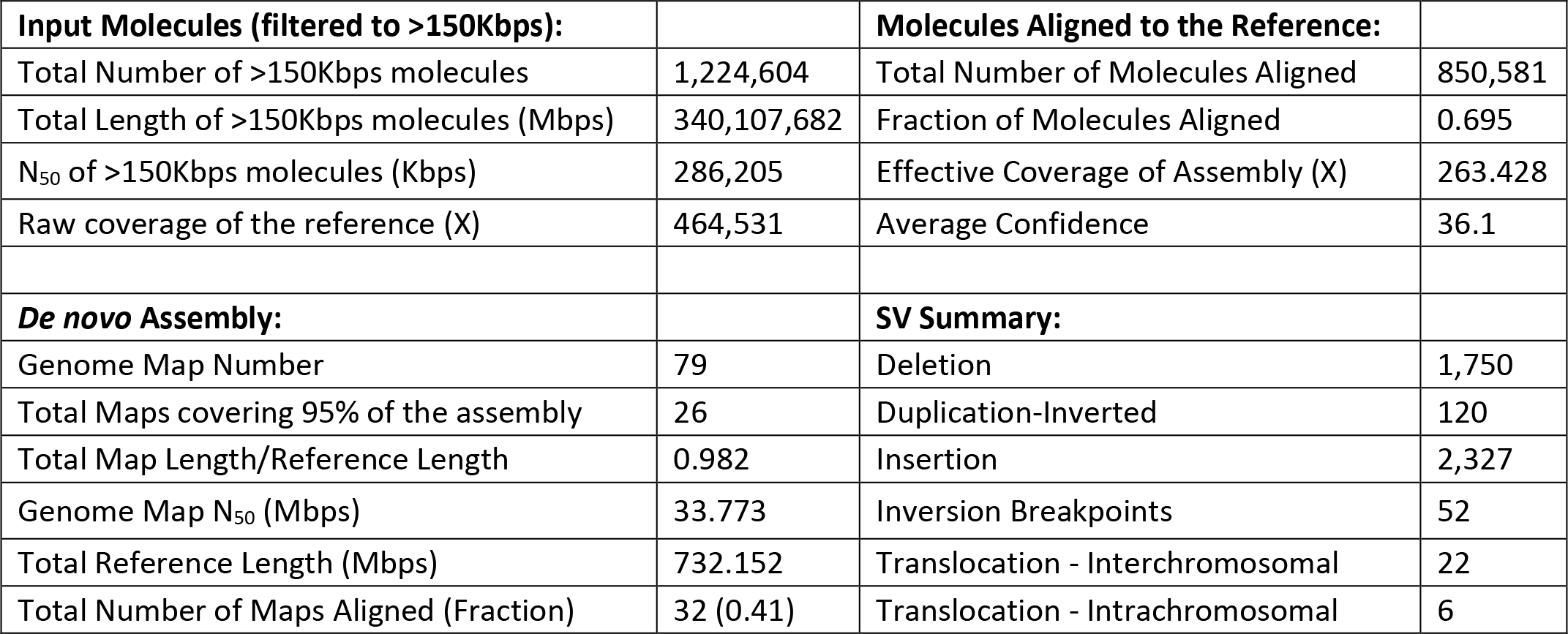
Summary of BioNano Genomics DLS optical map generation and assembly. Only molecules >150Kbps in length were assembled into contig maps. The resulting assembly was aligned against the public Sorghum bicolor v3.0.1 genome assembly (“Reference”) for comparison and detection of potential structural discrepancies.

| Input Molecules (filtered to >150Kbps): |  | Molecules Aligned to the Reference: |  |
| --- | --- | --- | --- |
| Total Number of >150Kbps molecules | 1,224,604 | Total Number of Molecules Aligned | 850,581 |
| Total Length of >150Kbps molecules (Mbps) | 340,107,682 | Fraction of Molecules Aligned | 0.695 |
| N<sub>50</sub> of >150Kbps molecules (Kbps) | 286,205 | Effective Coverage of Assembly (X) | 263.428 |
| Raw coverage of the reference (X) | 464,531 | Average Confidence | 36.1 |
| **De novo Assembly:** |  | **SV Summary:** |  |
| Genome Map Number | 79 | Deletion | 1,750 |
| Total Maps covering 95% of the assembly | 26 | Duplication-Inverted | 120 |
| Total Map Length/Reference Length | 0.982 | Insertion | 2,327 |
| Genome Map N<sub>50</sub> (Mbps) | 33.773 | Inversion Breakpoints | 52 |
| Total Reference Length (Mbps) | 732.152 | Translocation - Interchromosomal | 22 |
| Total Number of Maps Aligned (Fraction) | 32 (0.41) | Translocation - Intrachromosomal | 6 |

**Figure 2:**
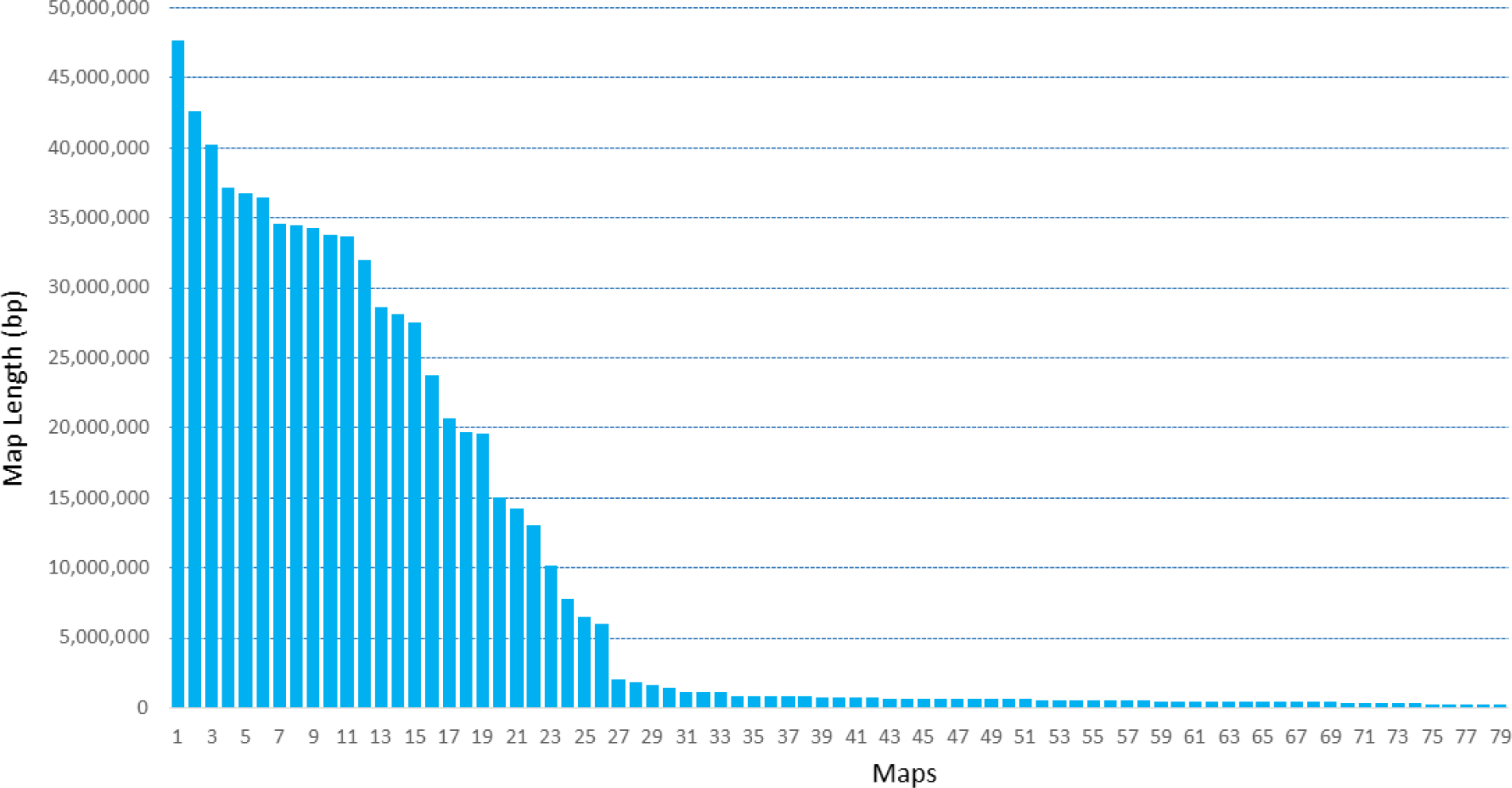
Summary of de novo assembly of single molecules into DLS optical maps. A total of 79 maps were created, out of which 32 accounts for 99.5% of the expected assembly size. The assembly N_50_ is 33.77Mbps while the largest map was 47.64Mbps in length.

The DLS optical maps were aligned to an *in-silico* map created from the reference *Sorghum bicolor* genome assembly to determine potential structural discrepancies between the two sorghum lines, Tx430 and BTx623 genome assemblies. With DLS, the majority of map breakages in plant inbred projects corresponds to long centromeric, ribosomal DNA and segmental duplication regions, as well as regions including residual structural heterozygosity. Results are summarized in Figure 3. In the end, all but three chromosomes were captured by only two maps, each one roughly corresponding to a chromosome arm. Only chromosome 5,9 and 10 provided more complexity and included 7, 5 and 4 maps, respectively. The alignments suggest the presence of substantial genomic rearrangements between the two maps. In total, near 52 inversion breakpoints were identified. For example, a large pericentromeric inversion, in relation to the optical map mapping to the same region, can be detected on chromosome 7 of BTx623 (Figure 3). Another large inversion is detected on chromosome 6 (Figure 3). Contig maps at the junctions exhibit robust individual molecule coverage, as shown for the left junction of the inversion on chromosome 6 (Supplementary Figure 2), suggesting the absence of any potential technical artifacts. Additional rearrangements include more than 3,000 insertion/deletion events and 28 translocations (Table 2). It remains to be determined whether some or all of those are related to assembly or mapping errors, or are actual structural variations between the two genotypes.

**Figure 3:**
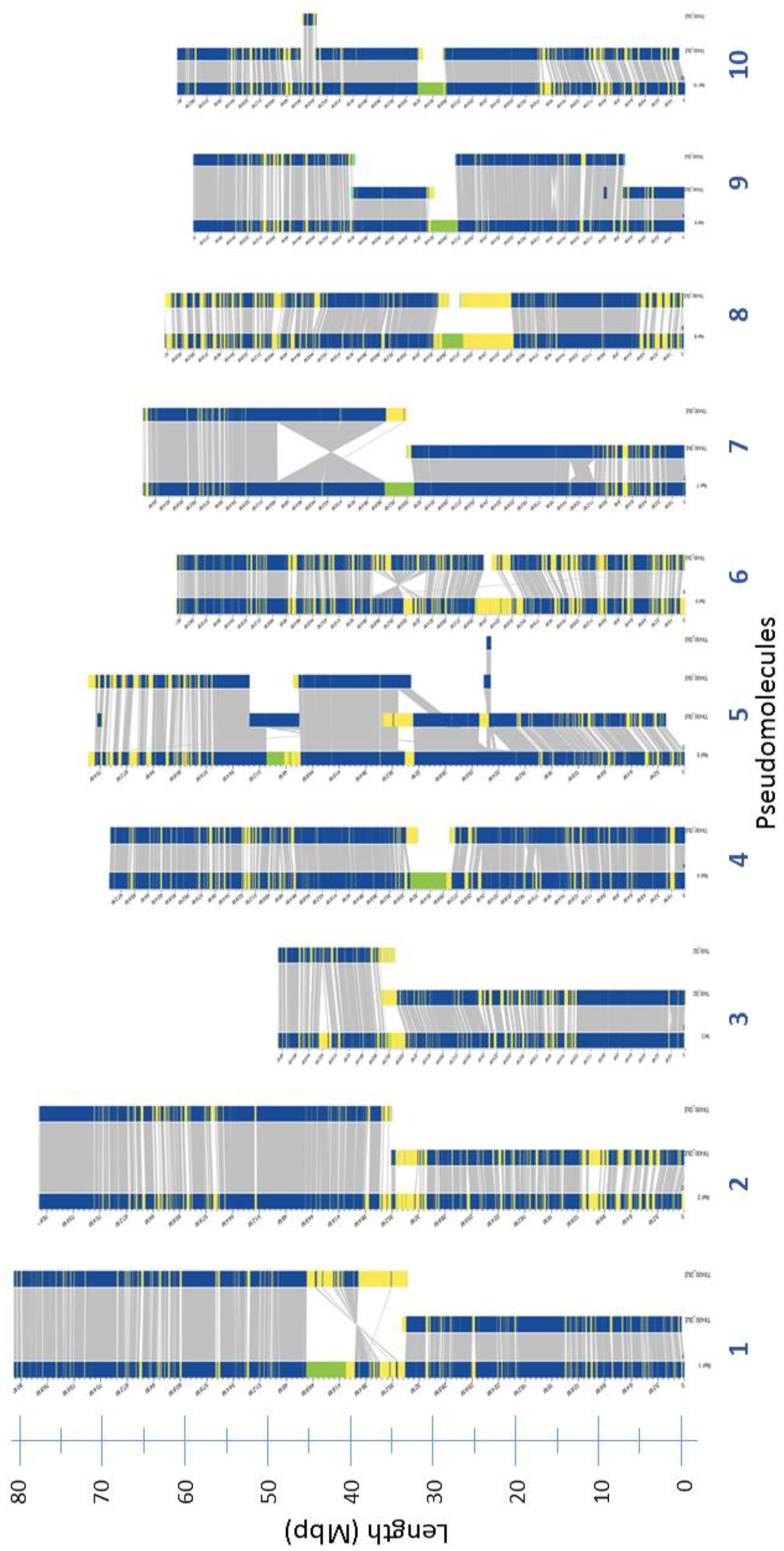
Alignment of DLS maps against in-silico maps of the public *Sorghum bicolor* reference assembly. Collinear DLE-1 markers on the two maps are linked (gray lines). All but three chromosomes are captured by two DLS maps. Large inversions on chromosomes 6 and 7 are marked by regions of reversed alignments between the two maps. Regions in green are stretches of random nucleotides in the reference assembly. Regions in yellow exhibit breaks in collinearity between the two maps.

### Generation of a chromosome-scale hybrid assembly

To further improve the assembly and determine the presence of chimeric contigs, the 724 contigs generated from the Pilon-polished SMARTdenovo assembly were scaffolded with the sorghum Tx430 DLS contig maps. The resulting hybrid scaffolds showed a significant improvement in contiguity when compared to the ONT contig assembly alone. The initial hybrid map assembly was made of 30 scaffolds, totaling 661.06Mbps, or 98.4% of the ONT public assembly, with a resulting N_50_ increasing from 3Mbps to 33.35Mbps (Table 3). A total of 141 conflicts in 83 ONT contigs were corrected during the hybrid map assembly process. Further inspection of the hybrid maps shows that small insertions in ONT contigs (Supplementary Figure 3) were repaired during the hybrid map creation process. An example of a hybrid scaffold (Figure 4) shows ONT contigs ordered and oriented after mapping to a single DLE optical map on chromosome 6. Apparent sequence gaps in the hybrid scaffolds may correspond to small genomic regions lacking sufficient number of DLE-1 labels to merge small ONT contigs.

**Figure 4:**
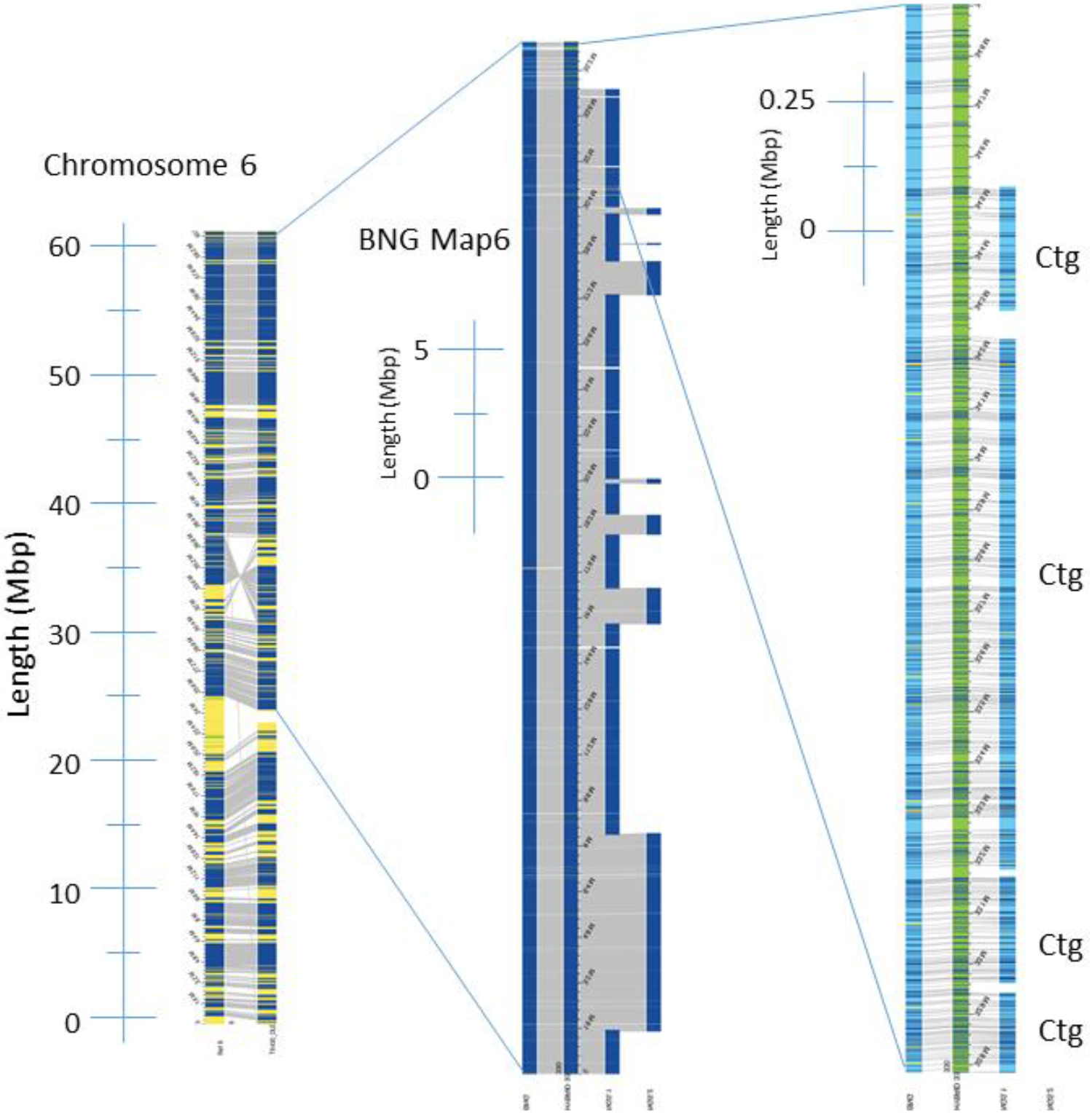
Detailed view of a region of Chromosome 6 and corresponding DLS maps. Left: DLS maps aligning to *in-silico* maps of Chromosome 6 from public v3.0.1 assembly; Center: Close-up view of DLS map 6 and mapped ONT contigs; Right: Close-up views of hybrid maps (center - green) generated by merging DLS maps (left - blue) and in-silico maps of ONT contigs (“Ctg”, right - blue). Distances are shown, in Mbps.

**Table 3:** Summary of DLS optical map assembly and merging with ONT contigs. Merging of DLS optical maps with ONT contigs produced a hybrid assembly made of 29 scaffolds, totaling 661.16Mbps, or >190% of the expected 732.1Mbps genome size, and with a N_50_ value of 33.28Mbps.

|  | Original<br>DLS<br>Genome<br>Map | Original<br>ONT<br>contigs | ONT<br>Contigs in<br>Hybrid<br>Scaffold | ONT Contigs<br>Not in<br>Hybrid<br>Scaffold | Hybrid<br>Scaffolds | Hybrid<br>Scaffolds<br>After<br>Correction |
| --- | --- | --- | --- | --- | --- | --- |
| Number<br>of Contigs | 79 | 723 | 500 | 363 | 30 | 29 |
| Total Length<br>(Mbps) | 719.339 | 671.867 | 644.44 | 25.117 | 661.06 | 661.16 |
| Minimum Contig<br>Length (Mbp) | 0.225 | 0.009 | 0.058 | 0.00006 | 0.086 | 0.115 |
| Maximum Contig<br>Length (Mbp) | 47.659 | 16.337 | 16.337 | 1.549 | 52.621 | 46.577 |
| N<sub>50</sub> Contig<br>Length (Mbp) | 33.773 | 3.053 | 2.991 | 0.13 | 33.35 | 33.28 |

The hybrid scaffolds were further assessed by comparing them with the Tx430 DLS contig maps previously aligned to the *Sorghum bicolor* v3.0.1 reference. The alignments indicated that the largest 52.62Mbps hybrid scaffold was chimeric, generated by merging two optical maps aligning to two different chromosomes from the public reference assembly. The chimera was created by a small ~40Kbps ONT contig sequence region with low DLE-1 marker count (Supplementary Figure 4), suggesting the presence of a conserved sequence. This 40Kb region was subsequently deleted from the ONT contig, splitting them in two separate contigs, before regenerating hybrid scaffolds with the same Tx430 contig maps. Correcting the chimeric scaffold did not significantly alter the overall hybrid map metrics. The corrected hybrid map assembly was made of 29 scaffolds, totaling ~661.16Mbps, with a slightly lower N_50_ length of 33.28Mbps (Table 3). To assess the completeness of the final hybrid map assembly, MUMmerplots were generated, following NUCmer 3.1 alignments of the 29 hybrid scaffolds described above to the *Sorghum bicolor* v3.0.1 genome assembly, and filtering for 1-on-1 alignments and rearrangements with a 20Kbps length cutoff. All 10 chromosomes from the public assembly were put in one unique plot (Figure 5). Results confirmed the presence of large inversions between the two genomes and suggested that the hybrid map assembly overall exhibits a high degree of completeness in relation to the public genome assembly.

**Figure 5:**
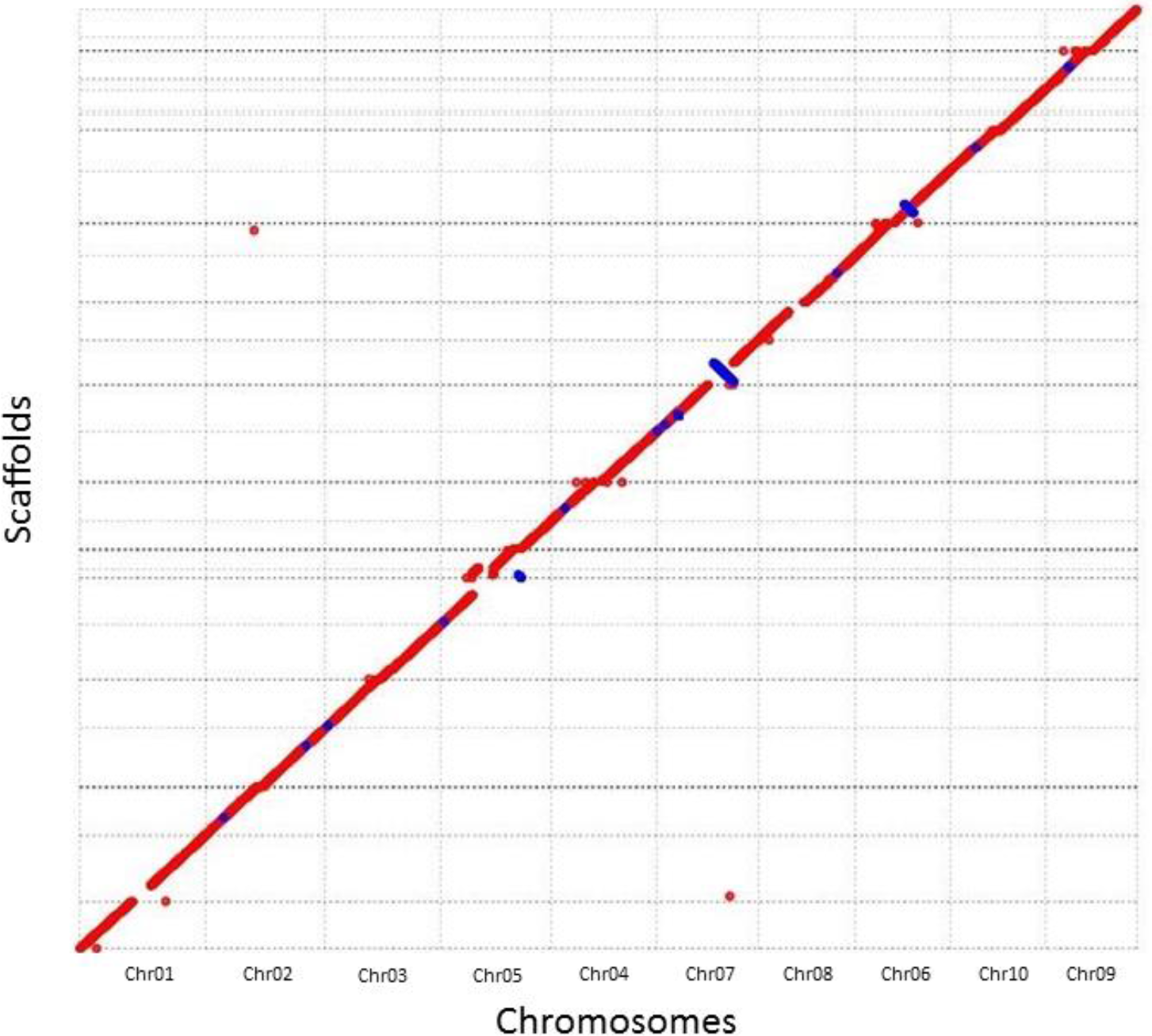
MUMmerplot comparison of Tx430 hybrid scaffolds (Y-axis) with the public *Sorghum bicolor* v3.0.1 genome assembly (X-axis). Hybrid scaffolds were aligned to all 10 BTx623 chromosomes using NUCmer and alignments were subsequently filtered for 1-on-1 alignment and rearrangements with a 20Kbps length cutoff. Chromosome order on the X-axis is related to the alignment of Tx430 scaffolds mapping to more than one chromosome. Chromosomal inversions and breakage in scaffold orientation in relation to the chromosomal sequence on the X-axis are shown as blue lines.

Finally, out of the 723 ONT contigs used to generate the hybrid scaffolds, 363 were not included, totaling approximately 25Mbps (Table 3). BLAST analysis of their sequences showed that 26 mapped to mitochondrial sequences and 97 mapped to plant pathogens. 208 out of the remaining 240 contigs showed best hits to sorghum sequences and a RepeatMasker analysis (26) of the 240 contigs indicated that 57.43% of their sequences were masked (including 51.88% matching with LTR elements present in Arabidopsis, rice, sorghum or maize). A separate tandem repeat analysis of the same 240 contigs with Tandem Repeat Finder (http://tandem.bu.edu/trf/trf409.linux64.download.html) shows that an additional 29.47% of sequences are masked (including two tandem repeats containing 3,913 and 2,403 copy numbers, respectively, of a 7bp motif). Taken together, 86.90% of the sequences present in those 240 contigs are masked. It is likely that most of them correspond to Tx430 conserved sequences mapping to existing scaffolds or to unmapped regions of the Tx430 genome.

## DISCUSSION

This study demonstrates that the Oxford Nanopore sequencing technology, combined with the DLS optical mapping technology recently developed by BioNano Genomics, can generate chromosome-scale assemblies of large and repeat-rich plant genomes. Assemblies of several large eukaryotic genomes with the Oxford Nanopore MinION sequencer have recently been attained in several studies. The ONT data presented here further demonstrate that the ONT-based assembly of the large and repeat-rich genome of sorghum exhibits a level of sequence completeness in the genome and, most importantly, in its genic regions, that is similar or higher to what has been observed in the public sorghum genome sequencing effort. It must be noted that ONT sequence accuracies described here are slightly lower than those observed in other nanopore-based eukaryotic genome assembly studies (6, 8). This could be explained by a combination of factors, including strategies or software tools used for base calling, correction and polishing, the overall nature, complexity and preponderance of conserved regions in the sorghum genome, the presence of residual heterozygosity linked to the fact that multiple Tx430 plants and seed sources were used for this project, or, in the case of comparisons performed against the public *Sorghum bicolor* v3.0.1 reference and annotation, structural variations between the two genotypes. In addition, Illumina whole-genome shotgun reads were used for polishing. It is therefore possible that Illumina polishing itself introduced low levels of sequencing errors, artificially decreasing the accuracy rate of the contig data.

The present study confirms that, while ONT sequences alone could generate an informative sorghum assemblies with high level of genome and genic completeness, the use of ultra-long-range genomics technologies such as BioNano Genomics optical maps can lead to chromosome-scale assemblies of a complex plant genome. DLS optical maps were used in the present study. Because labeling occurs directly without introducing nicks in the DNA strands, the average length and overall N_50_ of DLS optical maps tend to be significantly longer than the ones obtained using nicking enzymes, which further facilitated correcting, orienting and scaffolding of the ONT contigs. Alignments of the DLS optical maps generated in this study to the *Sorghum bicolor* reference genome showed that their sizes are mostly limited by the size of chromosomes. Several chimeric events were detected during merging of the ONT contigs with the DLS optical maps. This result is not unexpected and has been mentioned in other studies (11). It must be noted that, with sufficient sequencing coverage, ultra-long ONT reads (including reads over 1Mbp in length) may represent a useful complement to DLS optical maps. Similar N50 values as the one obtained here have been predicted in human when using ultra-long ONT sequencing reads at ~30X sequencing coverage (6).

The Tx430 assembly presented here can be further improved by manually editing the hybrid assembly and covering sequence gaps. Sequence accuracy also could be improved to reference-grade accuracy by increasing sequencing coverage or adding layers of sequencing information (10X Genomics linked reads, as an example, could be used to further polish the final assembly). The current state of the assembly however confirms that working-draft chromosome-scale scaffolds can be achieved rapidly, at reasonable costs, for large and complex plant genomes using existing long-read nanopore sequencing and DLS optical mapping technologies. These results have multiple implications. First, by greatly enhancing the contiguity of ONT assemblies, DLS optical maps are expected to greatly facilitate comparative genomics studies at the chromosomal level. Large pericentromeric inversions, such as the ones detected on chromosomes 6 and 7, could greatly impact marker detection and recombination in a breeding program involving parental lines distinguished structurally by such inversions. Contig sizes attained in this study also could facilitate the direct comparison of regions involved in specific traits of interest, such as disease-resistance gene clusters. Second, sequences in this study were generated using the USB-Sized MinION sequencer. Its overall portability means that informative whole-genome assemblies can now be attained in multiple settings and locations that, up until recently, didn’t have the means and resources to generate such assemblies. The *de novo* assembly of a complex plant genome also could be greatly accelerated by the adoption of larger ONT sequencing platforms, including the PromethION where up to 48 flow cells, each generating more than 100Gbps of data, can be run simultaneously in 48 (or more) hours. This could have a remarkable impact on how genomes are being determined and defined in the world of agricultural biotechnology, by allowing the rapid characterization, for example, of mutagenized lines or of transgene insertions, or by facilitating the design of multiple gene editing targets and allowing the direct prediction of potential off-target mutations in a large number of transformed plants.

## METHODS

### Plant material and genomic DNA extraction

Whole leaves were sampled from *Sorghum bicolor* accession Tx430 grown in the greenhouse for approximately two months. Samples were immediately collected in dry ice and stored at −80C until further handling. 25-30g of frozen tissue was ground in liquid nitrogen prior to extraction of high molecular weight genomic DNA from sorghum Tx430 adult leaf tissue using a protocol which was originally designed for the extraction of DNA for BAC library construction (18). For Bionano mapping, fresh tissue was collected from three-week-old seedlings. High molecular weight (HMW) genomic DNA was prepared according to a protocol developed specifically by BioNano Genomics. Briefly, 2g young leaf tissue was collected from live plants and immediately treated with a formaldehyde solution. After fixation, tissue was cut into approximately 2×2 mm squares and homogenized using a stator. Nuclei were isolated by gradient centrifugation and embedded into agarose plugs. After overnight proteinase K digestion in the presence of Lysis Buffer (BioNano Genomics) and 1 hour treatment with RNase A, plugs were washed three times in 1X Wash Buffer (BioNano Genomics), then five times in 1X TE buffer. After the final wash, DNA was recovered by melting the plugs, digesting agarose with 2μl of 0.5U/μl Agarase, and dialyzing for 45 minutes in 1X TE buffer.

### Oxford Nanopore MinION library construction and sequencing

For “short read” MinION runs, 2-3μg high molecular weight DNA was sheared to ~8Kbps with a g-Tube (Covaris) prior to library construction. For “long read” MinION runs, up to 100μl high molecular weight DNA at a concentration not exceeding 50ng/μl was size-selected to 20Kbps and above using a PippinHT system (Sage Science). The DNA recovered from the instrument was quantified and used directly for library construction.

“Short read” libraries and all but three “long read” libraries were built using the Ligation Sequencing Kit 1D (SQK-LSK108) from Oxford Nanopore. Briefly, DNA repair first was performed by incubating 2-4μg of genomic DNA using a FFPE Repair Mix enzymes (New England Biolabs) in 1X FFPE Repair Buffer for 15 minutes at 20°C, followed by cleanup with 0.5X AMPure XP beads and resuspension of the DNA in 45μl nuclease-free water. (For DNA samples with an initial DNA concentration lower than 45ng/μl, multiple DNA repair reaction, each containing 45μl of DNA, were performed and pooled together after repair and before bead cleanup.) End repair and dA-tailing were then performed simultaneously by mixing repaired DNA with 3μl Ultra II End-prep enzyme mix (New England Biolabs) in 1X Ultra II End-prep reaction buffer, and incubating 20 minutes at 20°C, followed by 20 minutes at 65°C. After cleanup with 0.5X AMPure XP beads, DNA was ligated with 1D Adapter Mix (Oxford Nanopore Technologies) in 1X Blunt/TA Ligation Mastermix (New England Biolabs) and incubated for 10 minutes at 20°C. Cleanup was performed by resuspending ligated DNA with 0.5X AMPure XP beads and washing the beads twice with 140μl Adapter Bead Binding buffer (Oxford Nanopore Technologies). After washing, DNA was dried down and resuspended in 12μl of Elution Buffer (Oxford Nanopore Technologies) before being combined with 35μl RBF buffer (Oxford Nanopore Technologies) and 28μl nuclease-free water and loaded on a MinION SpotON flow cell (FLO-MIN106). Sequencing was performed for 48 hours with a MinION Mk1B sequencer. The resulting FAST5 files were base called with v2.0.1 of the Albacore base caller. The remaining three “long read” libraries were prepared with the ONT Rapid Sequencing Kit 1D (SQK-RAD002). Briefly, 16μl of high molecular weight genomic DNA (corresponding to 2-4μg of DNA, less than the recommended 16μg of DNA) was mixed with 5μl FRN fragmentation mix (Oxford Nanopore Technologies) and incubated at 30°C for 1 minute, followed by 75°C for 1 minute. After incubation, 1μl Rapid 1D Adapter mix (Oxford Nanopore Technologies) and 1μl Blunt/TA Ligation Mastermix (New England Biolabs) were added and the library as incubated at 20°C for 30 minutes. Loading on a MinION flow cell (FLO-MIN106) was performed after mixing the library with 25.5μl RBF buffer (Oxford Nanopore Technologies) and 27.5μl nuclease-free water.

### Illumina sequencing

Illumina whole-genome shotgun sequencing was performed using the same high molecular weight DNA as the one used for MinION sequencing. Briefly, after randomly shearing ~1μg of DNA to ~600-700bps with a Covaris S2 Focused Ultrasonicator (Covaris), a whole-genome shotgun paired-end Illumina library was prepared using a KAPA Hyper Prep Kit (KAPA Biosystems), following the manufacturer’s recommendations, and sequenced on an Illumina HiSeq2500 to approximately 150X sequencing coverage equivalent of the sorghum genome. The resulting reads were trimmed and quality-filtered before polishing.

A 10X Genomics Chromium library was build using a separate high molecular weight DNA aliquot, following the manufacturer’s instruction, and sequenced on an Illumina HiSeq2500 to approximately 60X sequencing coverage of the sorghum genome. The resulting reads were subsequently trimmed and quality-filtered. Sequencing reads mapping to a region of the public v3.0.1 assembly containing the Sb-CENH3 gene, along with reads containing the same 10X barcode sequence as the mapped reads, were selected and assembled using the Supernova *de novo* assembler (10X Genomics).

### Oxford Nanopore MinION genome assembly and analysis

Oxford Nanopore MinION runs were base called using ONT Albacore Sequencing Pipeline Software (version 2.0.1) on the FAST5 files. Passed reads (Q score >= 7) of length 2K+bp from all MinION runs were fed into CANU (version 1.6) for read correction, followed by de novo assembly using SMARTdenovo (https://github.com/ruanjue/smartdenovo) on the corrected reads of length 5K+bp. CANU correction was run by setting corMaxEvidenceErate to 0.15 and correctedErrorRate to 0.12, while SMARTdenovo was run by setting the engine of overlapper to compressed kmer overlapper and the generate consensus to true. The SMARTdenovo consensus sequences were then polished with 100x Illumina reads using Pilon (version 1.21) for two iterations. RACON was also applied to contig polishing with Nanopore reads but was not used in the final assembly because the polishing did not improve the accuracy. Tandem repeats in non-scaffolded ONT contigs were detected using the Tandem Repeats Finder (http://tandem.bu.edu/trf/trf409.linux64.download.html).

The accuracy and completeness of polished contigs were evaluated by two methods. Full-length contigs were mapped to the public reference v3.0.1 using NUCmer 3.1 (MUMmer v3.9.4alpha) and the completeness comparison was performed by MUMmerplot 3.5 (MUMmer 3.23 package) on the NUCmer results after filtering to 1-on-1 alignments and allowing rearrangements with a 20Kbps length cutoff. Contigs were also chunked into 10Kbp fragments and mapped to the public reference using minimap2 (version 2.10) by allowing at most 3 secondary alignments. Average identity was calculated for the best hit of 50+% coverage for every chunk. To further evaluate the accuracy of polished contigs, Illumina reads were mapped to the contigs using BOWTIE2 and the average identity was calculated using uniquely mapped reads. Illumina reads were also mapped to the public reference v3.0.1 using BOWTIE2 for accuracy comparison. Lastly, the gene model set from the public reference v3.0.1 was mapped to the polished contigs using BOWTIE2 to evaluate the completeness of genic sequences coverage.

### BioNano Genomics DLS optical maps construction and hybrid assembly

Labeling and staining of the DNA were performed according to a protocol developed by BioNano Genomics. Labeling was performed by incubating 750ng genomic DNA with 1X DLE-1 Enzyme (BioNano Genomics) for 2 hours at 37°C, followed by 20 minutes at 70°C, in the presence of 1X DL-Green (BioNano Genomics) and 1X DLE-1 Buffer (BioNano Genomics). Following proteinase K digestion and DL-Green cleanup, the DNA was pre-stained by mixing the labeled DNA with 1X Flow Buffer (BioNano Genomics), in the presence of 1X DTT, and incubating overnight at 4°C. Staining was performed by adding 3.2μl of a DNA Stain solution (BioNano Genomics) for every 300ng of pre-stained DNA and incubating at room temperature for at least two hours before loading onto the BioNano Chip. Loading of the chip and running of the BioNano Genomics Saphyr System were all performed according to the Saphyr System User Guide (https://bionanogenomics.com/support-page/saphyr-system/). Data processing, construction of the DLS contigs and of the hybrid map assembly were all performed using the BioNano Genomics Access software suite.

### Data availability

All sequencing data that support the findings of this study have been deposited in the National Center for Biotechnology Information SRA database and are accessible through the SRA accession number SRP148505. All other relevant data are available from the corresponding authors on request.

## ACKNOWLEDGEMENTS

The authors wish to thank Ida Abbott and Praveena Parakkal for genomic DNA extraction, Matthew King for his scientific guidance, as well as Marissa Simon and Ping Che for their scientific guidance and for providing plant tissue for this project.

## AUTHOR CONTRIBUTIONS

S.D. and H.L. supervised the project; S.D., Y.Z., V.L. and H.L. conceived and designed the study; S.D. and V.L. performed the experiments; Y.Z., V.L., L.Y. and H.L. analyzed data; G.M. assisted with the conception and design of the study; S.D. and V.L. wrote the manuscript with input from all other authors.

## COMPETING INTERESTS

The authors declare no competing interests.

## MATERIALS & CORRESPONDENCE

Correspondence and requests for materials should be addressed to S.D or H.L.

